# Computational Geometric Tools for Modeling Inherent Variability in Animal Behavior

**DOI:** 10.1101/531392

**Authors:** Matthew T. Stamps, Soo Go, Ajay S. Mathuru

## Abstract

A fundamental challenge for behavioral neuroscientists is to represent inherent variability among animals accurately without compromising the ability to quantify differences between conditions. We developed two new methods that apply curve and shape alignment techniques to address this issue. As a proof-of-concept we applied these methods to compare normal or alarmed behavior in pairs of medaka (*Oryzias latipes*). The curve alignment method we call Behavioral Distortion Distance (BDD) revealed that alarmed fish display less predictable swimming over time, even if individuals incorporate the same action patterns like immobility, sudden changes in swimming trajectory, or changing their position in the water column. The Conformal Spatiotemporal Distance (CSD) technique on the other hand revealed that, in spite of the unpredictability, alarmed individuals share an overall swim pattern, possibly accounting for the widely held notion of “stereotypy” in alarm responses. More generally, we propose that these new applications of known computational geometric techniques are useful in combination to represent, compare, and quantify complex behaviors consisting of common action patterns that differ in duration, sequence, or frequency.

## 1 Introduction

Quantitative description of the behavior of an animal is a key element of neuroscience. As important as this is, even the most comprehensive ethograms fail to capture the inherent variability in the behaviors of individuals, whether they consist of simple or complex sets of actions [1, 2, 3, 4]. This is further compounded by the error-prone nature of subjective assessment during human observations. A pertinent issue for behavioral studies, therefore, is the development of appropriate methods for the quantitative comparison of behaviors between two animals or a single individual under two or more conditions.

Researchers typically quantify parameters (such as the average velocity or average height climbed) based on expert knowledge of the ethology of that species or the kinematics of a behavior. These measures are useful, yet they can be inadequate as descriptors of behavior for a variety of reasons. One notable reason is that they are gross approximations of the behavioral action being measured. Averaging of observational data may not be representative of the kinematics and can be confounded by what has been described elsewhere as “the failure of averaging” [2]. Furthermore, there can be considerable variability in the execution of even those behaviors that are considered “stereotypical” or instinctive.

As elaborated at great length by Schleidt in 1974, individual variability can exist in the form of “incompleteness of the elements” that make up an innate behavior or in the form of changes in the characteristics of those “elements” such as their duration, sequencing, or frequency within a given episode of executing the behavior [5]. Consequently, the above mentioned gross approximations are at best an imperfect representation of the behavior that neither capture nor do justice to the intra and inter subject variability [6]. Another reason measures fail to be adequate descriptors of behavior is that even if specific parameters are defined as relevant in a behavioral readout, they may not be available when examining a new species or when studying previously uncategorized behaviors. Quantification of infrequent events (such as episodes of immobility or jumps) is also somewhat unsatisfactory as a measure since it omits and obscures a large portion of the behavioral data. In the ideal scenario, an entire behavioral episode would be represented in as complete a form as possible during quantification.

Studies over the past few years have started to address these (or related) issues by applying tools from computational geometry, computer vision, and machine learning to describe behaviors [7]. These methods have proved useful, for example in examining the entire behavioral repertoire of the hydra [8], for automated annotation of behavior in fruit flies [9], identifying the temporal features that explain spontaneous swimming behavior in worms [10], identifying the dynamics of shoaling in fish [11], and detecting sub-second modules in mice behavior [12]. Such studies are providing new insights like the continuity between behavioral states [13] and objective means for comparison of new datasets to perform spatiotemporal mapping of posture of non-stereotyped actions [6].

Here, we report the development of two new methods for the representation and quantitative analysis of locomotion of pairs of animals moving freely in a two dimensional space. We applied these methods to compare swimming behavior of pairs of fish. The first among these methods is based on an approach to automated curve alignment developed by [14] that can be efficiently calculated with a computational technique called *dynamic time warping* [15]. Dynamic time warping has been applied in the past to analyze biological phenomena, such as speech patterns [16] and cardiac filling phases [17]. In recent years, others have employed techniques based on dynamic time warping to examine and compare temporal patterns and acoustic features in bottle-nosed dolphins [18], whales [19], and songbirds [20]. The same technique has also been adapted to compare locomotion and infer postures in humans [21, 22]. Motivated by a classic result in differential geometry called the Frenet-Serret Formula [23], we introduced the notion a *behavior curve* associated to an animal’s trajectory and derived a measure that we call the Behavioral Distortion Distance (BDD), which can be efficiently calculated via dynamic time warping to provide meaningful quantitative comparisons for entire episodes of locomotion.

The second method we developed examines the spatiotemporal kinetics encoded in heat maps. When comparing heat maps between animals that represent the time spent at each location in an arena, it is not useful to look at a straightforward average since hot spots from different animals will average out and disappear. Instead, one must first align the heat maps so that corresponding hot spots coincide in order to reveal if there are common patterns and similarities. We implemented a well-established algorithm [24, 25, 26, 27, 28] used previously in cortical cartography [29, 30], for finding conformal (angle preserving) alignments between surfaces - a heat map can be interpreted as a surface in 3-dimensional space where different intensities correspond to different heights in the z-direction - that uses discrete geometric objects called *circle packings*. We call the resulting measure between heat maps the Conformal Spatiotemporal Distance (CSD).

As a proof-of-concept, we asked if these methods can quantitatively distinguish normal swimming behavior from a behavior described as “alarmed” in fish. Though the species where such a phenomenon has been described have expanded considerably [31, 32, 33] since its discovery in minnows approximately 80 years ago [34], alarm behavior to conspecific injury has been mainly studied in the Ostariophysi species of fish [35, 36]. Even among Ostariophysi, species differ in the expression of alarmed behavior. Many different parameters, in captivity and in the wild, have been described for different fish including a change in inter-individual distance in shoals, freezing, darting, change in the vertical position in a water body, etc. [37, 38, 39, 40, 41]. As such, it is difficult to predict what an alarmed behavior will look like in a species where this has not been described before. We therefore chose as our test case the single published study from one of the authors (at the time we started this project) describing an alarm response in medaka [42]. We asked if the individuals in the two conditions (alarmed and normal) were distinguishable without an experimenter’s subjective criteria in defining what action sequence or pattern constitutes an alarmed state.

We found that individuals in each condition cluster together. Unexpectedly, the BDD between most alarmed individuals were also high. This suggests that swimming trajectories can become so unpredictable when medaka are alarmed that they may not match a second episode or a second individual expressing the same behavior. On the other hand, the CSD between most alarmed individuals were low, highlighting the existence of a discernible pattern in spite of low predictability. Overall, our results suggest that BDD and CSD together are useful aides to represent long episodes of swimming behavior. Our results also indicate that these computational geometric methods can be applied more universally for quantitative analysis of locomotion behavior of any kind, in any animal.

## 2 Results

### 2.1 Individual Variability in Swimming Patterns

We used a data-set that recorded the swimming motion of thirty-six medaka over a 10 minute interval [42]. The observational tanks were featureless small rectangular tanks with dimensions 20 cm × 12 cm × 5 cm (*L* × *H* × *W*). While the tanks restricted movement compared to the animals’ natural habitat, they allowed multiple degrees of freedom in swimming motion and can be accepted as reasonably “free” spaces for motion. In particular, the fish could swim in any manner across the tank approximately six to seven body lengths long, four body lengths deep, and one and a half body lengths wide. Half of the fish were exposed to an alarm substance *(Schreckstoff* or SS) at the same time point (the 2 minute mark) in the assay as depicted in (Figure 1A; SS) while the other half were exposed to a control substance (tank water; NSS). We used an automated tracker to record each animal’s position in the tank (Supplementary Video S1) and developed a measure of difference in behavior, or the BDD between two trajectories over a common length of time. Our model takes into account kinematic factors, such as the velocity (rate of displacement) of an animal at a given moment in time, as well as intrinsic geometric properties of the trajectory, such as curvature (the rate of change of turning while traveling at unit speed; see Figure 5 and the Methods section for details). Additionally, we included the vertical height in the tank as a parameter in our model because it is known to change for alarmed individuals in other species [43, 44, 45, 32, 37]. For all of the results listed in this article, BDD refers to the behavioral distortion distance indexed by height, velocity, and curvature, or BDD_Θ_ for Θ = {*y, v, κ*} in the notation of the Methods section.

**Figure 1:**
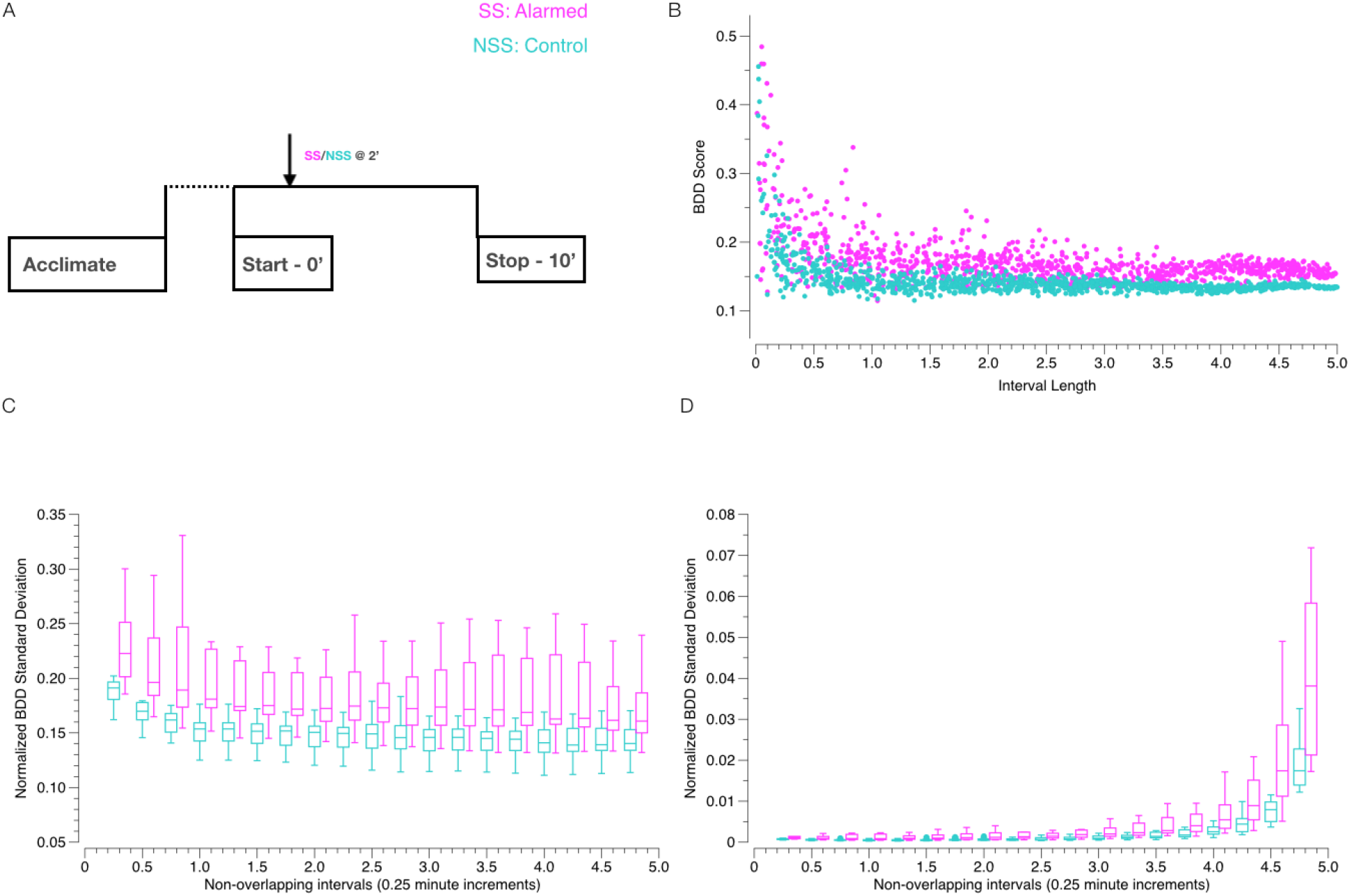
(A) Assay design: Alarm substance (SS; magenta) or tank water (NSS; cyan) was delivered into the assay chamber for 30 seconds at the 2 minute mark; (B) Intra-individual BDD over 1000 randomly generated pairs of nonoverlapping intervals of various lengths for a control (cyan) and an alarmed individual (magenta); (C) Box-and-whisker plots of mean intra-individual BDD for all control (cyan) and alarmed (magenta) fish over non-overlapping intervals with lengths between 0.25 and 4.75 minutes at 0.25 minute increments; (D) Box-and-whisker plots of normalized standard deviations of intra-individual BDD for all control (cyan) and alarmed (magenta) fish over non-overlapping intervals with lengths between 0.25 and 4.75 minutes at 0.25 minute increments.

To evaluate the differences in behavior between distinct individuals, we first set out to understand the amount of variability to expect within a single individual’s behavior over different periods of time. In particular, we sought to identify (1) the average BDD from an individual to itself at (non-overlapping) time intervals and (2) the length of time at which we see the least amount of variation in BDD from an individual to itself. An answer to (1) would provide a reasonable benchmark for the BDD between different individuals and an answer to (2) would suggest the time length over which we could obtain the most consistent/reliable BDD between different individuals.

As a preliminary step, we calculated the BDD from each fish to itself over a thousand randomly generated pairs of non-overlapping intervals of various lengths between 0 and 5 minutes. Figure 1B shows an example of the BDD values calculated for a representative alarmed (SS) or non-alarmed (NSS) fish. Each point in the plot corresponds to a pair of intervals with the length of the intervals plotted along the x-axis and their corresponding BDD along the y-axis. The calculations revealed that the BDD values of both the alarmed and non-alarmed fish exhibited significant variation for shorter periods of time, but quickly stabilized as the length of the time intervals increased. This is expected since the set of pairs of non-overlapping intervals with longer duration is smaller than the set of such intervals with shorter duration. For instance, there is only one non-overlapping pair of intervals of length 5 minutes, so only one BDD per fish of this duration can be generated. The stability we observed, however, made us optimistic that the questions raised by (1) and (2) could be answered precisely.

We ran the following, more structured, computation: For each fish in the experiment and interval length d between 0.25 and 4.75 minutes at 0.25 minute increments, we randomly generated a hundred pairs of non-overlapping time intervals of length *d* and calculated the BDD for the fish over each pair of intervals. We then calculated the mean and standard deviation of the values for each fish and choice of *d*. Since the size of the set of pairs of non-overlapping intervals of length *d* varies as a function of *d*^2^, we normalized the standard deviations of the BDD for each *d* by the size of the corresponding domain. The mean and normalized standard deviations (standard deviation of BDD per unit area of input) are shown in Table 3 with corresponding box plots in Figure 1C.

**Table 1:** The means and standard deviations of BDD for pairs of fish over 30 second intervals, broken down into three types: pairs of control fish (NSS-NSS), pairs of alarmed fish (SS-SS), and mixed pairs with one control and one alarmed fish (NSS-SS). The “Distribution” columns refer to the distribution of alarmed fish in the lower and upper half of fish ordered by mean BDD to all other fish in the experiment (Split) and corresponding likelihood of such a split if the fish had been ordered randomly.

| Duration<br>(minutes) | Mean BDD |  |  | Standard Deviation |  |  | Distribution |  |
| --- | --- | --- | --- | --- | --- | --- | --- | --- |
|  | NSS-NSS | NSS-SS | SS-SS | NSS-NSS | NSS-SS | SS-SS | Split | Likelihood |
| 0-0.5 min | 0.182 | 0.183 | 0.189 | 0.022 | 0.028 | 0.028 | 56-44 | 74% |
| 0.5-1 min | 0.187 | 0.179 | 0.178 | 0.026 | 0.024 | 0.025 | 50-50 | 100% |
| 1-1.5 min | 0.174 | 0.177 | 0.179 | 0.020 | 0.024 | 0.026 | 50-50 | 100% |
| 1.5-2 min | 0.181 | 0.179 | 0.179 | 0.023 | 0.021 | 0.021 | 44-56 | 74% |
| 2-2.5 min | 0.184 | 0.187 | 0.182 | 0.031 | 0.028 | 0.025 | 50-50 | 100% |
| 2.5-3 min | 0.182 | 0.198 | 0.213 | 0.029 | 0.032 | 0.043 | 28-72 | 1.8% |
| 3-3.5 min | 0.170 | 0.206 | 0.222 | 0.017 | 0.037 | 0.048 | 11-89 | 0.00052% |
| 3.5-4 min | 0.181 | 0.206 | 0.222 | 0.024 | 0.041 | 0.049 | 28-72 | 1.8% |
| 4-4.5 min | 0.171 | 0.226 | 0.262 | 0.018 | 0.099 | 0.127 | 28-72 | 1.8% |
| 4.5-5 min | 0.186 | 0.209 | 0.222 | 0.026 | 0.036 | 0.044 | 33-67 | 9.4% |
| 5-5.5 min | 0.175 | 0.257 | 0.309 | 0.019 | 0.132 | 0.165 | 17-83 | 0.015% |
| 5.5-6 min | 0.173 | 0.252 | 0.302 | 0.019 | 0.129 | 0.160 | 17-83 | 0.015% |
| 6-6.5 min | 0.188 | 0.253 | 0.298 | 0.025 | 0.131 | 0.163 | 33-67 | 9.4% |
| 6.5-7 min | 0.186 | 0.254 | 0.305 | 0.025 | 0.129 | 0.160 | 28-72 | 1.8% |
| 7-7.5 min | 0.184 | 0.247 | 0.293 | 0.025 | 0.130 | 0.166 | 39-61 | 32% |
| 7.5-8 min | 0.176 | 0.232 | 0.263 | 0.024 | 0.101 | 0.131 | 22-78 | 0.22% |
| 8-8.5 min | 0.181 | 0.224 | 0.250 | 0.024 | 0.100 | 0.133 | 33-67 | 9.4% |
| 8.5-9 min | 0.174 | 0.240 | 0.295 | 0.021 | 0.133 | 0.166 | 33-67 | 9.4% |
| 9-9.5 min | 0.182 | 0.245 | 0.297 | 0.028 | 0.133 | 0.165 | 33-67 | 9.4% |
| 9.5-10 min | 0.186 | 0.243 | 0.295 | 0.024 | 0.133 | 0.168 | 44-56 | 74% |

**Table 2:** The means and standard deviations of CSD for pairs of fish over 1 minute intervals, broken down into three types: pairs of control fish (NSS-NSS), pairs of alarmed fish (SS-SS), and mixed pairs with one control and one alarmed fish (NSS-SS). The “Distribution” columns refer to the distribution of alarmed fish in the lower and upper half of fish ordered by mean CSD to all other fish in the experiment (Split) and corresponding likelihood of such a split if the fish had been ordered randomly.

| Duration<br>(minutes) | Mean CSD |  |  | Standard Deviation |  |  | Distribution |  |
| --- | --- | --- | --- | --- | --- | --- | --- | --- |
|  | NSS-NSS | NSS-SS | SS-SS | NSS-NSS | NSS-SS | SS-SS | Split | Likelihood |
| 0-2 min | 0.013 | 0.014 | 0.014 | 0.0039 | 0.0040 | 0.0038 | 33-67 | 9.4% |
| 2-4 min | 0.014 | 0.013 | 0.011 | 0.0035 | 0.0037 | 0.0038 | 61-39 | 32% |
| 4-6 min | 0.014 | 0.012 | 0.010 | 0.0038 | 0.0036 | 0.0037 | 67-33 | 9.4% |
| 6-8 min | 0.014 | 0.012 | 0.010 | 0.0041 | 0.0040 | 0.0035 | 67-33 | 9.4% |
| 8-10 min | 0.014 | 0.012 | 0.010 | 0.0041 | 0.0036 | 0.0035 | 61-39 | 32% |
| 0-10 min | 0.015 | 0.013 | 0.011 | 0.0041 | 0.0040 | 0.0040 | 72-28 | 1.8% |

**Table 3:** The summary statistics for BDD over various time intervals between 0.25 and 3 minutes (with 0.25 minute increments), broken down into three types of pairs: pairs of control fish (NSS-NSS), pairs of alarmed fish (SS-SS), and mixed pairs with one control and one alarmed fish (NSS-SS), corresponding to Figure 1 C and D.

| Length | Mean |  |  | Std. Dev. |  |  | Normalized Std. Dev. |  |  |
| --- | --- | --- | --- | --- | --- | --- | --- | --- | --- |
|  | All | NSS | SS | All | NSS | SS | All | NSS | SS |
| 0.25 min | 0.213 | 0.187 | 0.238 | 0.054 | 0.033 | 0.076 | 0.0012 | 0.0007 | 0.0017 |
| 0.50 min | 0.192 | 0.168 | 0.216 | 0.041 | 0.022 | 0.059 | 0.0010 | 0.0006 | 0.0015 |
| 0.75 min | 0.187 | 0.160 | 0.214 | 0.037 | 0.018 | 0.056 | 0.0010 | 0.0005 | 0.0015 |
| 1.00 min | 0.182 | 0.155 | 0.210 | 0.034 | 0.015 | 0.053 | 0.0011 | 0.0005 | 0.0017 |
| 1.25 min | 0.178 | 0.153 | 0.203 | 0.032 | 0.013 | 0.051 | 0.0012 | 0.0005 | 0.0018 |
| 1.50 min | 0.176 | 0.150 | 0.201 | 0.031 | 0.012 | 0.050 | 0.0013 | 0.0005 | 0.0020 |
| 1.75 min | 0.175 | 0.150 | 0.200 | 0.029 | 0.012 | 0.046 | 0.0014 | 0.0006 | 0.0022 |
| 2.00 min | 0.173 | 0.148 | 0.199 | 0.027 | 0.011 | 0.043 | 0.0015 | 0.0006 | 0.0024 |
| 2.25 min | 0.172 | 0.147 | 0.198 | 0.026 | 0.011 | 0.042 | 0.0017 | 0.0007 | 0.0028 |
| 2.50 min | 0.172 | 0.147 | 0.197 | 0.020 | 0.010 | 0.030 | 0.0016 | 0.0008 | 0.0024 |
| 2.75 min | 0.172 | 0.146 | 0.197 | 0.020 | 0.009 | 0.031 | 0.0020 | 0.0009 | 0.0031 |
| 3.00 min | 0.173 | 0.146 | 0.200 | 0.018 | 0.008 | 0.027 | 0.0022 | 0.0010 | 0.0034 |
| 3.25 min | 0.174 | 0.146 | 0.201 | 0.016 | 0.008 | 0.025 | 0.0027 | 0.0013 | 0.0041 |
| 3.50 min | 0.173 | 0.145 | 0.200 | 0.017 | 0.007 | 0.026 | 0.0037 | 0.0016 | 0.0058 |
| 3.75 min | 0.172 | 0.144 | 0.199 | 0.015 | 0.007 | 0.024 | 0.0048 | 0.0021 | 0.0076 |
| 4.00 min | 0.170 | 0.144 | 0.197 | 0.013 | 0.006 | 0.021 | 0.0066 | 0.0028 | 0.0104 |
| 4.25 min | 0.169 | 0.144 | 0.195 | 0.012 | 0.005 | 0.020 | 0.0109 | 0.0043 | 0.0175 |
| 4.50 min | 0.168 | 0.144 | 0.192 | 0.010 | 0.004 | 0.017 | 0.0209 | 0.0079 | 0.0339 |
| 4.75 min | 0.168 | 0.145 | 0.192 | 0.007 | 0.003 | 0.011 | 0.0536 | 0.0209 | 0.0863 |

Table 3: The summary statistics for BDD over various time intervals between 0.25 and 3 minutes (with 0.25 minute increments), broken down into three types of pairs: pairs of control fish (NSS-NSS), pairs of alarmed fish (SS-SS), and mixed pairs with one control and one alarmed fish (NSS-SS), ...
| Length | Min |  | Q1 |  | Median |  | Q3 |  | Max |  |
| --- | --- | --- | --- | --- | --- | --- | --- | --- | --- | --- |
|  | NSS | SS | NSS | SS | NSS | SS | NSS | SS | NSS | SS |
| 0.25 min | 0.162 | 0.186 | 0.181 | 0.204 | 0.191 | 0.223 | 0.196 | 0.247 | 0.202 | 0.368 |
| 0.50 min | 0.146 | 0.165 | 0.162 | 0.186 | 0.170 | 0.196 | 0.177 | 0.231 | 0.179 | 0.409 |
| 0.75 min | 0.141 | 0.154 | 0.151 | 0.175 | 0.162 | 0.189 | 0.168 | 0.237 | 0.175 | 0.431 |
| 1.00 min | 0.130 | 0.152 | 0.146 | 0.173 | 0.157 | 0.181 | 0.163 | 0.220 | 0.178 | 0.415 |
| 1.25 min | 0.125 | 0.145 | 0.144 | 0.171 | 0.154 | 0.174 | 0.159 | 0.210 | 0.176 | 0.383 |
| 1.50 min | 0.125 | 0.145 | 0.141 | 0.167 | 0.152 | 0.175 | 0.158 | 0.202 | 0.172 | 0.385 |
| 1.75 min | 0.123 | 0.146 | 0.139 | 0.166 | 0.152 | 0.172 | 0.156 | 0.202 | 0.169 | 0.457 |
| 2.00 min | 0.120 | 0.142 | 0.138 | 0.162 | 0.150 | 0.172 | 0.156 | 0.199 | 0.171 | 0.452 |
| 2.25 min | 0.120 | 0.141 | 0.137 | 0.162 | 0.150 | 0.175 | 0.155 | 0.203 | 0.169 | 0.466 |
| 2.50 min | 0.116 | 0.138 | 0.137 | 0.160 | 0.149 | 0.173 | 0.157 | 0.193 | 0.179 | 0.540 |
| 2.75 min | 0.114 | 0.137 | 0.136 | 0.157 | 0.146 | 0.172 | 0.155 | 0.199 | 0.183 | 0.526 |
| 3.00 min | 0.115 | 0.136 | 0.134 | 0.157 | 0.146 | 0.174 | 0.153 | 0.207 | 0.184 | 0.536 |
| 3.25 min | 0.115 | 0.134 | 0.135 | 0.158 | 0.146 | 0.171 | 0.153 | 0.214 | 0.187 | 0.552 |
| 3.50 min | 0.114 | 0.132 | 0.134 | 0.157 | 0.145 | 0.171 | 0.151 | 0.221 | 0.189 | 0.521 |
| 3.75 min | 0.113 | 0.132 | 0.134 | 0.157 | 0.144 | 0.169 | 0.150 | 0.217 | 0.190 | 0.536 |
| 4.00 min | 0.111 | 0.132 | 0.133 | 0.158 | 0.141 | 0.163 | 0.152 | 0.213 | 0.192 | 0.526 |
| 4.25 min | 0.112 | 0.133 | 0.133 | 0.156 | 0.139 | 0.163 | 0.153 | 0.208 | 0.195 | 0.513 |
| 4.50 min | 0.113 | 0.133 | 0.134 | 0.153 | 0.139 | 0.162 | 0.153 | 0.192 | 0.199 | 0.532 |
| 4.75 min | 0.114 | 0.132 | 0.134 | 0.150 | 0.140 | 0.161 | 0.152 | 0.185 | 0.200 | 0.564 |

The results of this computation show that the mean BDD for the control group (NSS) is fairly level for *d* ≥ 0.5 after an initial substantial decline from *d* = 0.25. This indicates that the BDD from a control fish to itself over durations of 30 seconds or more is consistent. On the other hand, the normalized standard deviations of BDD for the control group (NSS) has a slight “J” shape with an initial decline from *d* = 0.25 followed by a rapidly increasing incline for *d* ≥ 1 with minimum between *d* = 0.5 and *d* = 0.75. This indicates that the least variation in BDD for individuals occurs over 30 to 45 second intervals, which means we can expect to obtain the most consistent comparisons of BDD over such durations.

This analysis also revealed that the mean BDD values for alarmed fish were significantly higher than that of non-alarmed fish (Figure 1C) for the majority of lengths of comparison. As seen in Figure 1C, the first quartile of BDD values for alarmed fish is greater than the third quartile of the BDD values for non-alarmed fish for every duration except *d* = 4.75. In other words, more than 75% of the BDD values for fish exposed to the alarm substance were strictly greater than at least 75% of the BDD values for fish exposed to control substance. This suggests that it is possible to predict whether or not a fish can be considered alarmed with a specified level of confidence by looking only at its BDD values with *itself*, that is, without the need to compare it to any other fish [46].

### 2.2 Behavioral Distortion Distances (BDD) between Alarmed and Non-Alarmed Medaka

With the least variation in BDD for individuals occurring over 30 second durations, we calculated the BDD between each pair of the thirty-six medaka over 30 second intervals throughout the 10 minute experiment. The results are shown in Figure 2 and detailed in Table 1.

**Figure 2:**
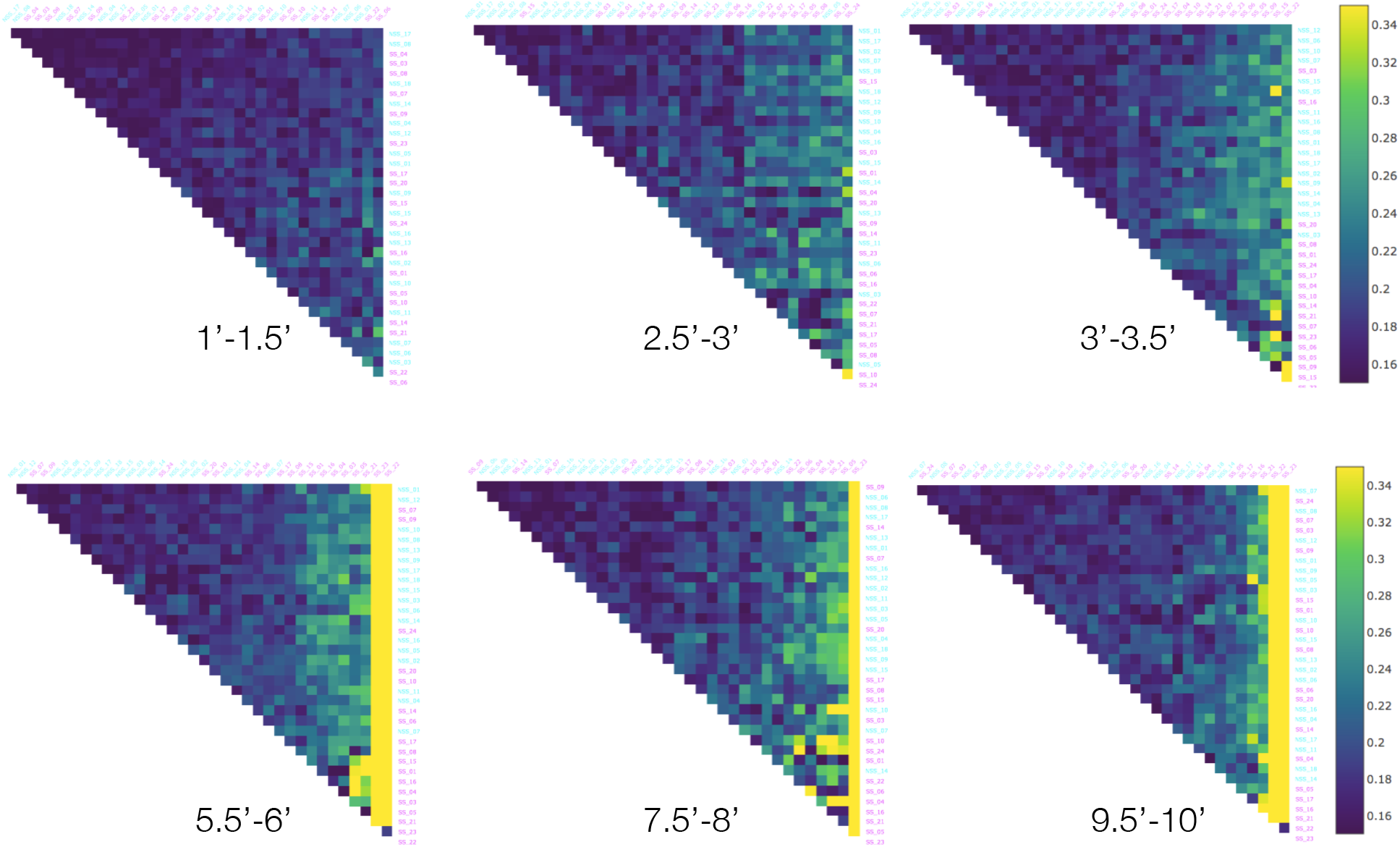
The Behavorial Distortion Distances (BDD) between all pairs of the thirty-six fish in the dataset over selected 1 minute intervals. Alarmed (SS; magenta) and control (NSS; cyan) individuals are ordered from lowest to highest centrality, i.e., average BDD to all other fish in the experiment.

As seen in the Table 1, the mean BDD within the control fish (NSS-NSS) was fairly constant throughout the experiment ranging between 0.173 and 0.187, only slightly higher than the mean BDD of 0.168 we observed in the intra-individual comparisons. The mean BDD between alarmed fish (SS-SS) and between the control and alarmed fish (NSS-SS) were also comparable to the control fish, but only for the first two minutes. They exhibit a significant increase after the 2 minute mark. We interpret this to be the consequence of exposure to the alarm substance delivered at the 2 minute mark as the conditions for each group were identical up to the that point. A similar pattern appears in the standard deviations of BDD, except that the values do not exhibit a significant jump until after the 4 minute mark.

In addition, we calculated the average BDD from each fish to all the other fish over each of the time intervals and sorted the fish from lowest to highest for each interval (Figure 2). For the first 2 minutes, the two groups (NSS and SS) were evenly mixed with respect to this ordering. As time progressed, the two groups became more and more distinguished with most of the control fish at the lower end of the spectrum and most alarmed at the higher end. This pattern was most apparent during the 3-3.5 minute interval. Table 1 shows the *distribution split*, that is, the percentage of SS fish among the lower half in the ordering versus the percentage in the upper half of the order, and the corresponding likelihood of such a split if the fish had been ordered at random. Between 3 and 3.5 minutes, the likelihood of obtaining the ordering we found is on the order of 1 in 200000. This suggests that fish exposed to alarm substance have a distinct swimming behaviour a short while after exposure to *Schreckstoff*, but not immediately.

### 2.3 Conformal Spatiotemporal Distances (CSD) between Alarmed and Non-Alarmed Medaka

We conducted a similar analysis for the Conformal Spatiotemporal Distance (CSD) method we developed to compare heat maps representing the amount of time a fish spends in each location of the observational tank throughout the experiment or a specified portion of it (see Figure 7 and Methods). We noted that in order for discernible patterns to emerge in these heatmaps, longer intervals than those used for BDD were required. We used 2 minutes intervals over the course of the 10 minute experiment along with the entire 10 minute interval(Figure 3) and detailed in Table 2.

**Figure 3:**
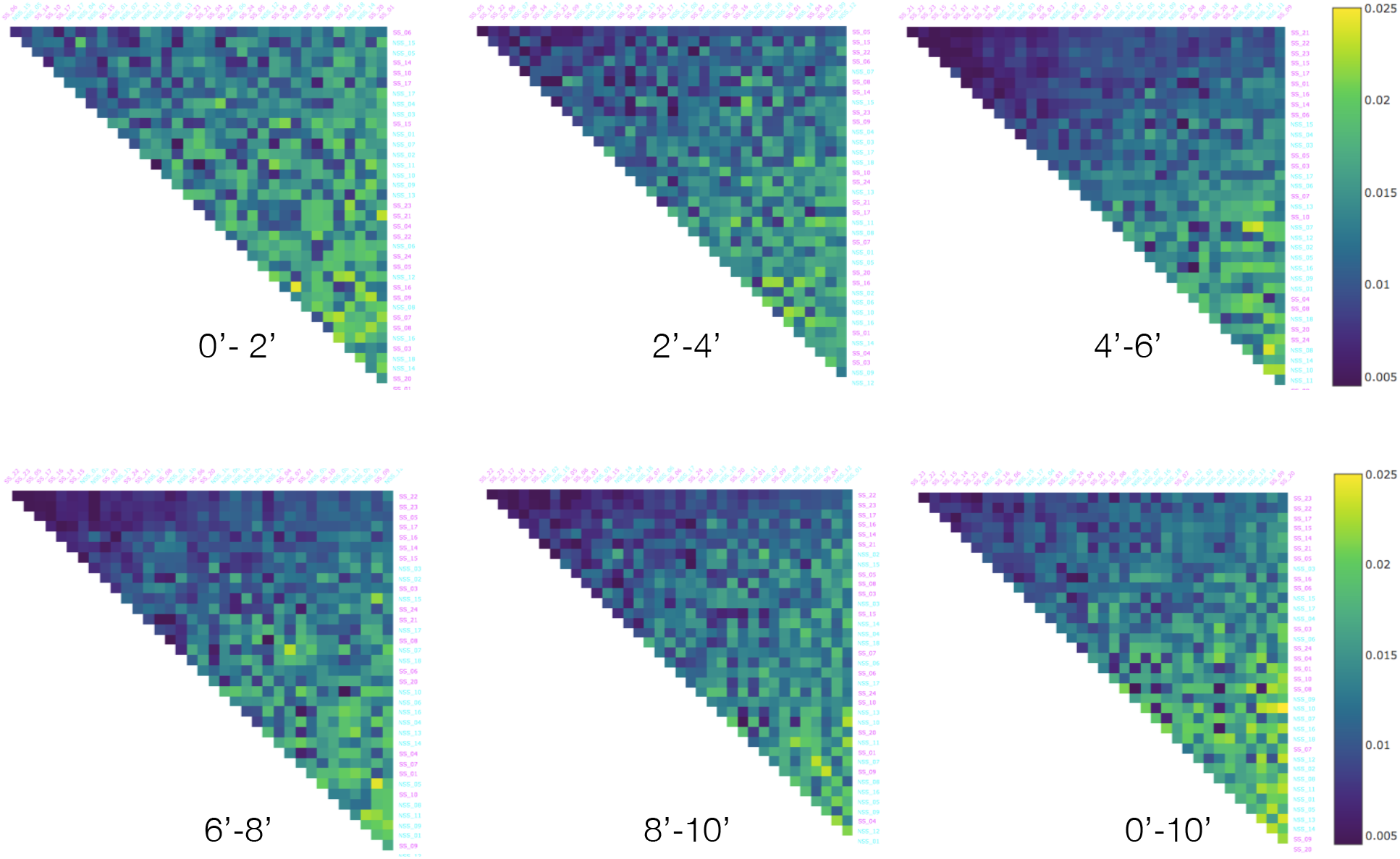
The Conformal Spatiotemporal Distances (CSD) between all pairs of the thirty-six fish in the dataset over selected 2 minute intervals as well as the full 10 minute interval. Alarmed (SS; magenta) and control (NSS; cyan) individuals are ordered from lowest to highest centrality, i.e., average CSD to all other fish in the experiment.

The mean CSD within the control fish (NSS-NSS) was essentially constant over the five 2 minute intervals ranging between 0.022 and 0.023. Once again, like the BDD,the mean difference between alarmed fish (SS-SS) and between the control and alarmed fish (NSS-SS) were comparable to control fish for the first 2 minutes, but exhibited a noticeable change after this point. However, unlike the BDD, there was drop in CSD for alarmed fish. The most substantial drop occurred during the 4-6 minute interval. The standard deviations did not exhibit any clear changes for any of the three comparisons.

As with BDD, we calculated the average CSD from each fish to all the other fish over each of the time intervals and sorted the fish from lowest to highest for each interval Figure 3. Initially, there were twice as many control fish than alarmed in the first half of this ordering, but as time progressed, that trend reversed with most of the alarmed at the lower end of the spectrum. This pattern was most apparent during the 6-8 minute interval and was mirrored by the entire experiment (0-10 minute). However, as Table 2 shows the likelihood of obtaining the ordering we found for the entire interval duration was less than 1 in 50, we consider CSD less capable of identifying a “typical” alarmed behavior. While the CSD did not perform as well as BDD when it came to classifying the state, it did capture a level of similarity in the behavior between alarmed fish that was not captured by BDD.

## 3 Discussion

Complex behaviors with common action patterns or modules, albeit in exaggerated or in a compressed forms, can be precluded from identification as different. We designed two new applications of known techniques in computational geometry for curve and surface alignment that are particularly relevant when examining complex behavior and behavioral states. We applied these techniques to reexamine the report of the existence of an alarm response in medaka [42]. Gross parameters such as the total duration of immobility among fish exposed to the alarm substance or *Schreckstoff* were adequate to differentiate two groups of medaka in the original report, even though individuals showed considerable variability. The analyses presented in this paper provide methods to quantify behaviors that are intrinsically variable and confirm that medaka behavior changes after exposure to *Schreckstoff* derived from conspecifics. This can be inferred without expert knowledge of the underlying ethology.

This report also provides some new insights into the alarm behavior of medaka that may be generalizable to other fish. First, the BDD analysis revealed that fish exposed to *Schreckstoff* differed from each other almost as much as they differed from non-alarmed fish. This may appear counter-intuitive at first glance if we expect the alarmed fish to behave in a comparable manner or more “uniformly” when expressing fear, however, these differences in BDD are consistent with the idea that normal kinematics are interrupted when medaka are alarmed. As a consequence, the repetitive aspects in their swimming are reduced and the swimming trajectories become less predictable overall. This is likely to be advantageous as an anti-predator strategy. Second, this analysis also revealed that the BDD from a fish to itself (Figure 1) can be a good indicator of its behavioral state. However, the caveat here is that one would need a length of recording of normal behavior (either from the same animal or comparable individuals) to establish the magnitudes of small and large BDD values.

One question that often arises in such analyses is not just whether two episodes are different, but how exactly do they differ. Imortantly, we can use our model to examine which parameters of the swimming kinematics contribute most to the difference between two behavoiral episodes. We cannot only determine whethor or not the behaviors of two animals were different for a given episode, but we can identify and measure *how* they were different. For instance, the differences in BDD throughout our results are dominated by curvature only about 5 - 10% of the time, while velocity dominates 40 - 45% of the time, and vertical height in the tank dominates the remaining time for the first 2 minutes across all animals and for control fish (NSS-NSS pairs) over the whole 10 minutes. After the 2 minute mark, however, there is a noticeable drop in velocity domination for most alarmed fish (SS-SS pairs; (Figure 4) and a corresponding increase in the domination of parameter vertical height in the tank. When comparing control fish against alarmed (most SS-NSS pairs) however, velocity domination increases with a corresponding decrease in the relevance of vertical height (not shown). This analysis can be interpreted most congruously in the following manner - that most alarmed medaka are fairly similar to each other when it comes to velocity changes, in particular periods of immobility, but they differ from each other primarily because of the location in the water column where they become immobile. In contrast, they differ from control fish in the frequency and the duration of immobility rather than the position occupied in the water column.

**Figure 4:**
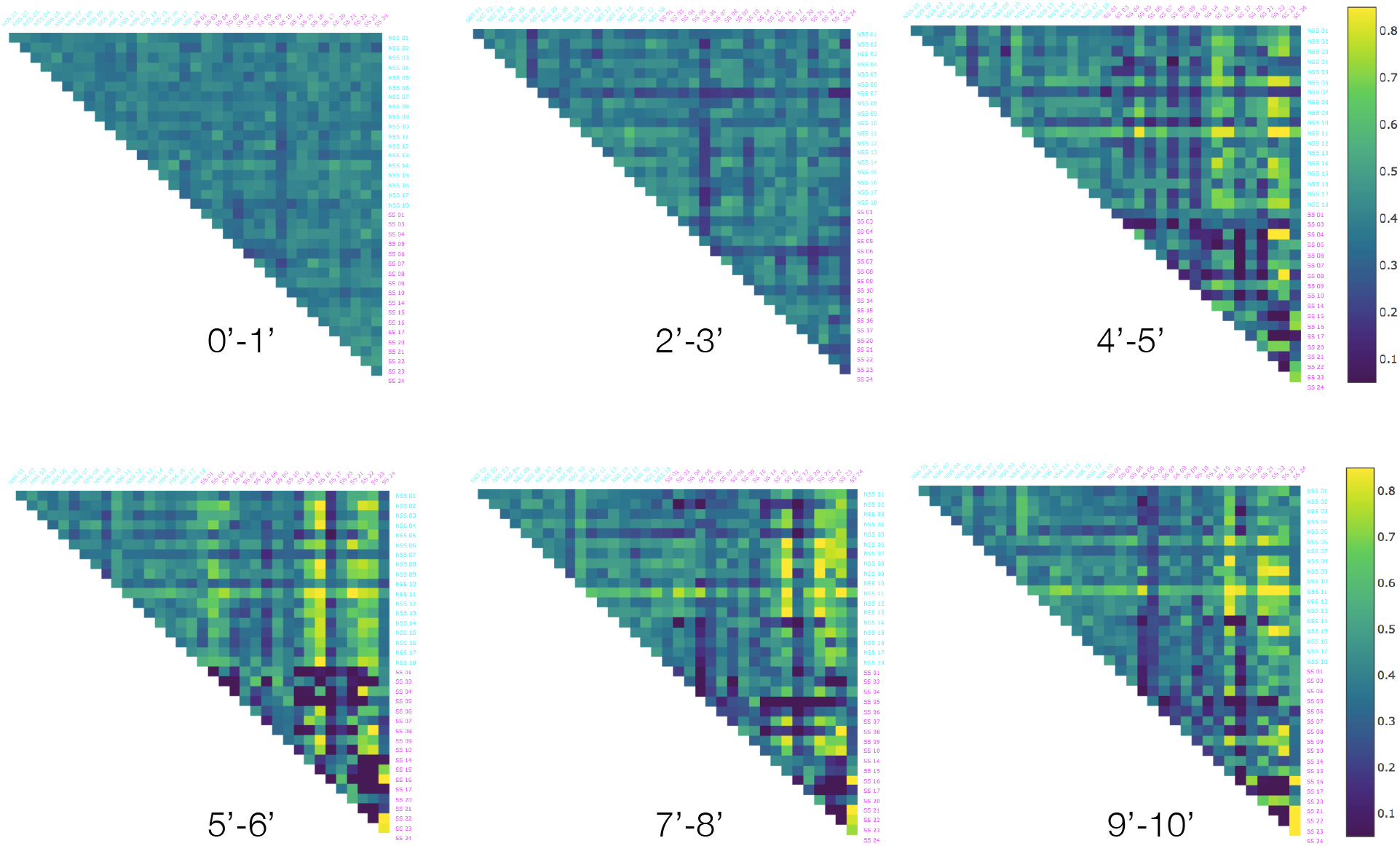
The percentage of time velocity was most significant factor (greatest contributor) instead of height and curvature in the calculation of BDD over a selection of 1 minute intervals. Starting at the 4 minute mark, velocity is the dominant factor for the majority of the time in each interval for most of the mixed alarmed/normal (NSS-SS) pairs, while it is the dominant factor for only a small portion of the time in each interval for most of the alarmed (SS-SS) pairs.

The CSD analysis was less effective than BDD for classification; however, it revealed a possible reason for human observers describing alarm behavior as being stereotypic in spite of considerable variability between individuals expressing alarm. As described in [5], “observer’s eye and mind perform a complex task of averaging and assigning weights to many different characteristics”, essentially integrating information over a large experimental time period. The CSD shows that most alarmed fish (60%; Figure 3) share an overall pattern of similarity which might be picked up by the observers as features more frequent in alarmed fish. Our analyses also reveal that control fish can exhibit “an alarmed state” and that alarmed state is an extension of the normal behavior “phase space”.

A general caveat of using the BDD and CSD for analysis is that they can only be applied to a pair of curves (resp. heat maps) at a time. It therefore accompanies a degree to difficulty in visualizing a characteristic feature, if any, common to all individuals in a condition when comparing two conditions. As demonstrated here, it is possible to overcome this limitation to an extent by statistical analysis of all pairs. We expect these methods would be able to verify a proposed model for “typical” behavior, however, by showing that the BDD or CSD between the candidate model and a sufficiently large sample of control animals were less than or equal to the intra-individual BDD (resp. CSD) over different time intervals. Another caveat of these measures is the assumption that the behavior being analyzed can be described by the intrinsic geometric and kinematic properties of locomotion. Indeed, behavioural posture is unaccounted in these analyses. However, here this limitation exists because of the recording technique employed. It can be overcome if postural data is identified in videos with higher resolution recordings that could then be incorporated to compute the BDD.

More generally, we have developed the curve alignment process so that it can be used to compare any two curves that change over a common length of time. We have also devised algorithms that generate parametric curves from tracking data normally obtained for analysis of behavior (that is, *x*- and *y*-coordinates, time-stamp, frame rate, etc.) These new applications can therefore be used to compare locomotion dependent behavior of any animal tracked in a two dimensional arena. It can also be applied to animals moving in three dimensional arena with simple modifications. We expect these methods will be more widely used after this demonstration of their utility. Accordingly, our software will be publicly available on a GitHub repository once this manuscript has been accepted for publication. In the meantime, please contact one of the authors for an up-to-date version of our python package.

## 4 Methods

### 4.1 Behavioral data

The data used for the development of methods comes from [42]. Details of the experimental setup, including the method for preparation of the alarm substance (*Schreckstoff*), can be found in the methods section of the original publication. The original tracking data used in our model is uploaded to public data repository and is provided with this paper.

### 4.2 Curve Alignment & Behavioral Distortion Distance (BDD)

Our first method compares the behavior of two animals by comparing the intrinsic geometric properties and dynamics of their trajectories.

#### 4.2.1 Model

We assume that the trajectory of each fish is a continuous, smooth parametric curve given by a vector-valued function **r**: [*t*_0_, *t_f_*] → ℝ^3^ where *t*_0_ and *t_f_* denote the start and end times of a specified time interval, respectively. To identify the geometric and dynamic features of the trajectories that are most relevant to behavior, we first calculate the rate at which an animal’s motion is changing relative to its current frame of reference. We model an animal’s frame of reference at a point along its trajectory with its *Frenet-Serret frame* at that point: For a non-degenerate, continuously differentiable, parametric curve *C* with the vector-valued function **r**: [*t*_0_, *t_f_*] → ℝ^3^ it is well known in differential geometry [23] that the unit tangent, normal, binormal vectors, defined by

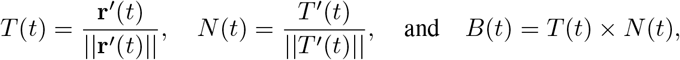

respectively, at each point of the trajectory form an orthonormal basis of ℝ^3^. Here, *g′*(*t*) denotes the derivative of a function *g* with respect to *t*, ║v║ denotes the magnitude of a vector *v* ∈ ℝ^3^, and × denotes the cross product. In otherwords, we can recoordinatize ℝ^3^ according to a fish’s position and orientation at each moment in time. Measuring the rate of change of an animal’s frame of reference at a given point is equivalent to calculating the rate of change of its Frenet-Serret frame, which is given by the famous Frenet-Serret formula, which we state below as a theorem.

##### Theorem 1

(Frenet-Serret Formula, [47, 48]). *If* **r**: [*t*_0_, *t_f_*] → ℝ^3^ *is a non-degenerate, continuously differentiable, parametric curve and T*(*t*), *N*(*t*), *and B*(*t*) *are its unit tangent, normal, and binormal vectors at time t, then*

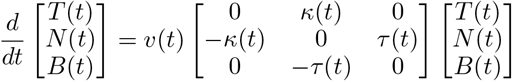

*where v*(*t*) = ║**r**(*t*)║, 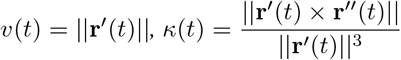, *and* 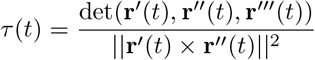.

The upshot of the Frenet-Serret formula in our context is that the rate of change of the Frenet-Serret frame is completely determined by *v*(*t*), *κ*(*t*), and *τ*(*t*), which are the *speed, curvature*, and *torsion*, respectively, of **r** at time *t*. While speed will be familiar to many readers as the rate of displacement, the curvature indicates the rate at which the curve is turning while traveling at unit speed, and the torsion indicates the rate which the curve is twisting while traveling at unit speed. Portions of the trajectory, velocity, and curvature of the Medaka NSS 01 fish over the first thirty seconds of its experiment are shown in Figure 5.

**Figure 5:**
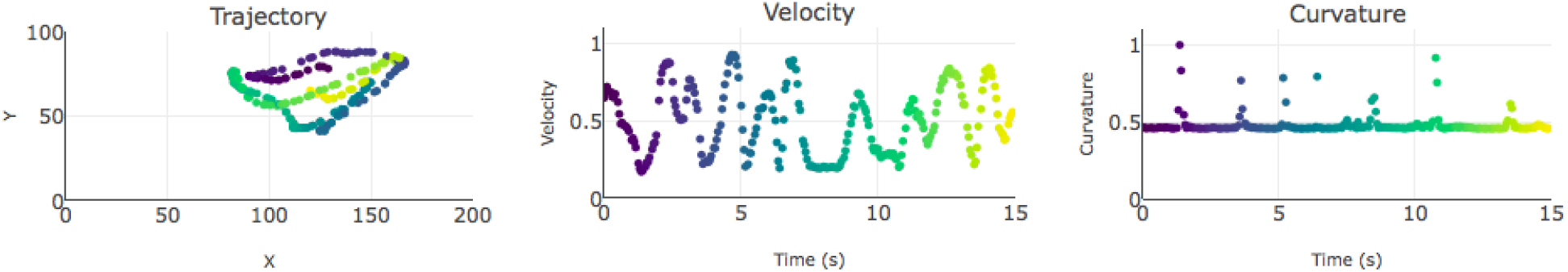
The trajectory, velocity, and curvature plots of the first 15 seconds of the fish NSS 01. The same color scale is used in each plot so corresponding points on each curve are colored with the same color.

We also considered the height of the fish in the tank as a behavioral factor, since it has been observed in other species that alarmed fish tend to dive more deeply than unalarmed fish. We did not, however, take into account the horizontal position in the tank, since left/right preferences and orientations have not been previously associated meaningfully in fish in the context of alarm behavior. For example, we consider two fish swimming at constant rates in circles of the same radius at constant heights in the tank to exhibit the same behavior even if one is swimming clockwise and the other counter-clockwise when viewed from above.

Now that we have identified several appropriate factors for comparing behavior, we must determine how to measure the difference between those factors. To do this, we introduce the following notion:

##### Definition 2.

Let **r**: [*t*_0_, *t_f_*] → ℝ^3^ be the trajectory of an animal and let Θ = {*θ*_1_,…, *θ_m_*} be a set of *m* behavioral factors, such as *v, κ*, and *τ*. The *behavior curve* of the trajectory indexed by Θ is the parametric curve *C*_Θ_: [*t*_0_, *t_f_*] → ℝ^m^ given by *C*_Θ_(*t*) = (*θ*_1_(*t*),…, *θ_m_*(*t*)).

Because it is virtually impossible to deliver an external stimulus simultaneously to two freely swimming animals (the stimulus takes time to disperse through the tank), it is not meaningful to compare two behavior curves at common moments in time. For instance, a slight time lag due to different reaction times could yield arbitrarily large quantitative differences between two animals that exhibit identical sequences of behaviors with that approach. Instead, we must identify an appropriate correspondence between pairs of points from each behavior curve in order to make a meaningful comparison. This is a curve alignment problem, which we address using a method developed by [14].

##### Definition 3.

An *alignment* between two parametric curves *C*^1^: [*t*_0_, *t_f_*] → ℝ^*d*_1_^ and 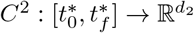 is a function

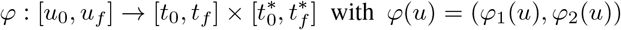

satisfying the conditions that 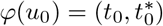, 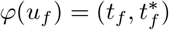, and *φ_i_* is monotone increasing and differentiable for each *i* ∈ {1, 2}. Such a function associates a unique point on *C*^2^ to each point on *C*^1^ by matching *C*^1^(*t*) to *C*^2^(*t*^*^) where 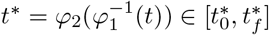 for each *t* ∈ [*t*_0_, *t_f_*].

An example alignment between the behavior curves with Θ = {*v*} for the first 30 seconds of observation for the Medaka NSS 01 fish and Medaka SS 01 fish is illustrated in Figure 6.

For every alignment *φ* between a fixed pair of parametric curves *C*^1^ and *C*^2^, we can calculate the amount of energy required to bend, stretch, and/or compress *C*^1^ onto *C*^2^ according to *φ*.

**Figure 6:**
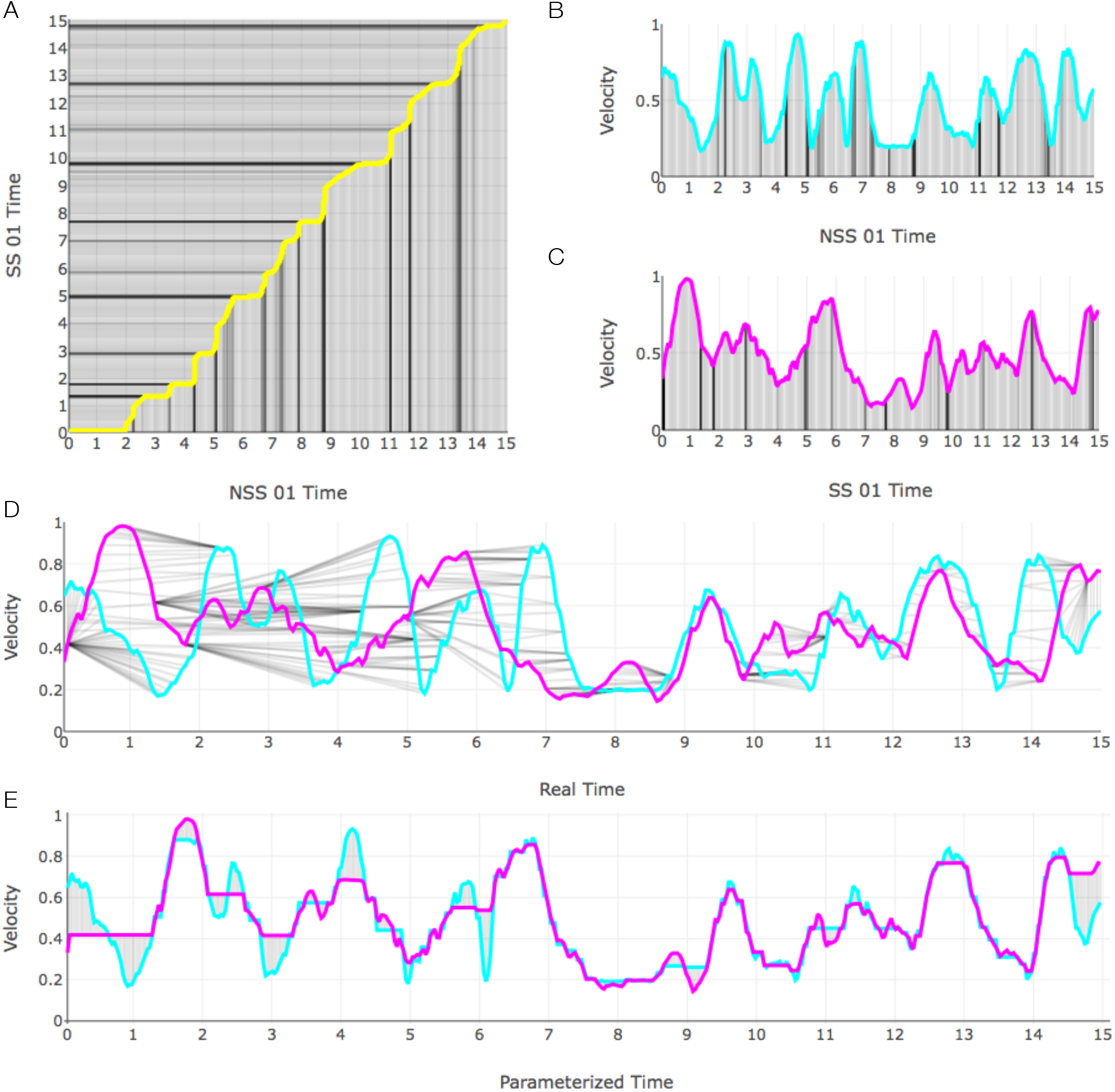
An alignment between the velocity curves of the NSS 01 and SS 01 medaka over the first 15 seconds of the experiment. (A) The optimal alignment *φ*_min_; (B) the velocity curve for NSS 01; (C) the velocity curve for SS 01; (D) the two curves plotted simultaneously with pairs of matched points indicated by gray line segments; (E) the two curves plotted with parameterized or “warped” time.

##### Definition 4.

Let *C*^1^: [*t*_0_, *t_f_*] → ℝ^*d*^ and 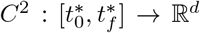 be parametric curves in *d*-dimensional space. The *distortion energy* of an alignment 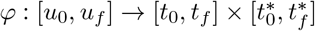 between *C*^1^ and *C*^2^ is

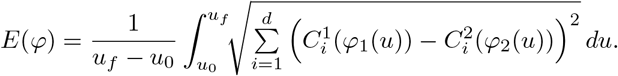

To compare the behaviors between two trajectories, we search for an alignment between their corresponding behavior curves that minimizes the distortion energy.

##### Definition 5.

Let **r**_1_: [*t*_0_, *t_f_*] → ℝ^*d*^ and 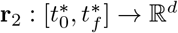 be the trajectories of two animals and let Θ = {*θ*_1_,…, *θ_m_*} be a set of behavioral factors. The *behavioral distortion distance* between **r**_1_ and **r**_2_ with respect to Θ is

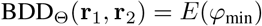

where *φ*_min_ is an alignment between the behavior curves 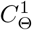 and 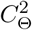 corresponding to **r**_1_ and **r**_2_, respectively, with minimum distortion energy.

The BDD is a modification of the *edit distance, d*_edit_, between parametric curves presented in [14], which they show to be a *pseudometric*, that is, (1) *d*_edit_(*C*^1^, *C*^1^) = 0, (2) *d*_edit_(*C*^1^, *C*^2^) = *d*_edit_(*C*^2^, *C*^1^), and (3) *d*_edit_(*C*^1^, *C*^3^) ≤ *d*_edit_(*C*^1^, *C*^2^) + *d*_edit_(*C*^2^, *C*^3^) for all parametric curves *C*^1^, *C*^2^, and *C*^3^ in a common Euclidean space. The fact that BDD is a pseudometric follows from very similar arguments. The edit distance and BDD fail to be metrics, however, since it is possible for distinct trajectories to have (edit or behavioral distortion) distance 0. This is perfectly fine in our context because it is reasonable for animals with distinct trajectories to exhibit identical behaviors.

#### 4.2.2 Implementation

We obtain the trajectories from the “track objects” algorithm in the MetaMorph software as sequences of *x*- and *y*-coordinates in the real plane. While the observational tank is 3-dimensional, the camera only captures the projection of a 2-dimensional, 20 cm × 12 cm, vertical cross-section of an animals movement. As described in [42], since videos were recorded from the front of the tank, displacement in the orthogonal width could not be recorded, effectively reducing a three dimensional space to a two dimensional observation. Ideally, we would be able to capture 3-dimensional trajectories; however, the BDD has the same derivation for 2-dimensional trajectories. The Frenet-Serret frames of a plane curve are given by the tangent and normal vectors and the Frenet-Serret formula is identical with torsion constantly equal to 0.

To approximate BDD, we use *dynamic time warping* (DTW) as proposed by [14]. Our implementation of the DTW algorithm is based on the version described in [49] with the distance function modified to account for velocity, curvature, and height. In particuar, we extended the existing DTW method within the Machine Learning module (mlpy) in Python to accommodate our use of multivariate data and different choices of Θ since the original DTW function only accepts two univariate sequences as data. While our method is able to handle input data of an arbitrarily high dimension (3-dimensional data and any number of behavioral factors), in our analyses thus far, we have only used *x*-coordinates, *y*-coordinates, velocity, and curvature. The velocity and curvature values are generated from the *x*- and *y*-coordinates using the formulas

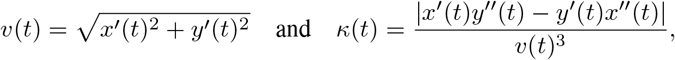

where the derivatives are computed by applying the gradient function in the NumPy package in Python to the *x* and *y* time series. In addition, we apply a simple moving average smoothing function to the first derivatives *x′*(*t*) and *y′*(*t*) before computing the velocity and curvature values in order to reduce the data noise. In our tests, the smoothing window is 10 frames, spanning 5 frames prior up to 4 frames following, for instance,

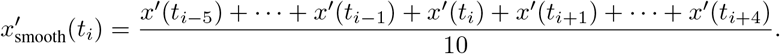

Furthermore, to account for variations in average values of behavioral factors between animals and differences in magnitudes between behavioral factors themselves, we normalize all of the data using the sigmoid function

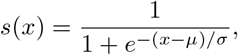

where *μ* and *σ* are the mean and the standard deviation of the respective input sequence. Since the range of the sigmoid function is [0,1], the value of BDD_Θ_ is bounded above by the number of factors in Θ.

### 4.3 Surface Alignment & Conformal Spatiotemporal Distance (CSD)

The second method we developed compares the spatiotemporal distributions between two animals. When comparing heat maps of time spent at each location in an arena between animals, it is not useful to look at a straightforward average since hot spots from different animals will average out and disappear. Instead, one must first align the heat maps so corresponding hot spots coincide in order to reveal if there are common patterns and similarities.

#### 4.3.1 Model

To examine the overall spatiotemporal patterns of an animal within an enclosure, we study its spatiotemporal distribution function, that is, the continuous function *ρ*: Ω → ℝ where Ω is the enclosure of the animal and *ρ*(*z*) is the number of seconds the animal spends at location *z* within Ω over a specified time interval *I*. When Ω is a region in the plane, this function can be visualized/interpreted as a heatmap where small values of *ρ* correspond to “cold” points and large values of ρ correspond to “hot” points. Alternatively, the spatiotemporal distribution function can be visualized/interpreted as the surface in ℝ^3^ with points {(*x, y, ρ*(*x, y*) | (*x, y*) ∈ Ω}, as illustrated in Figure 7. We assume that this surface is smooth and Riemannian, that is, it is equipped with a well-defined notion of distance and angles.

**Figure 7:**
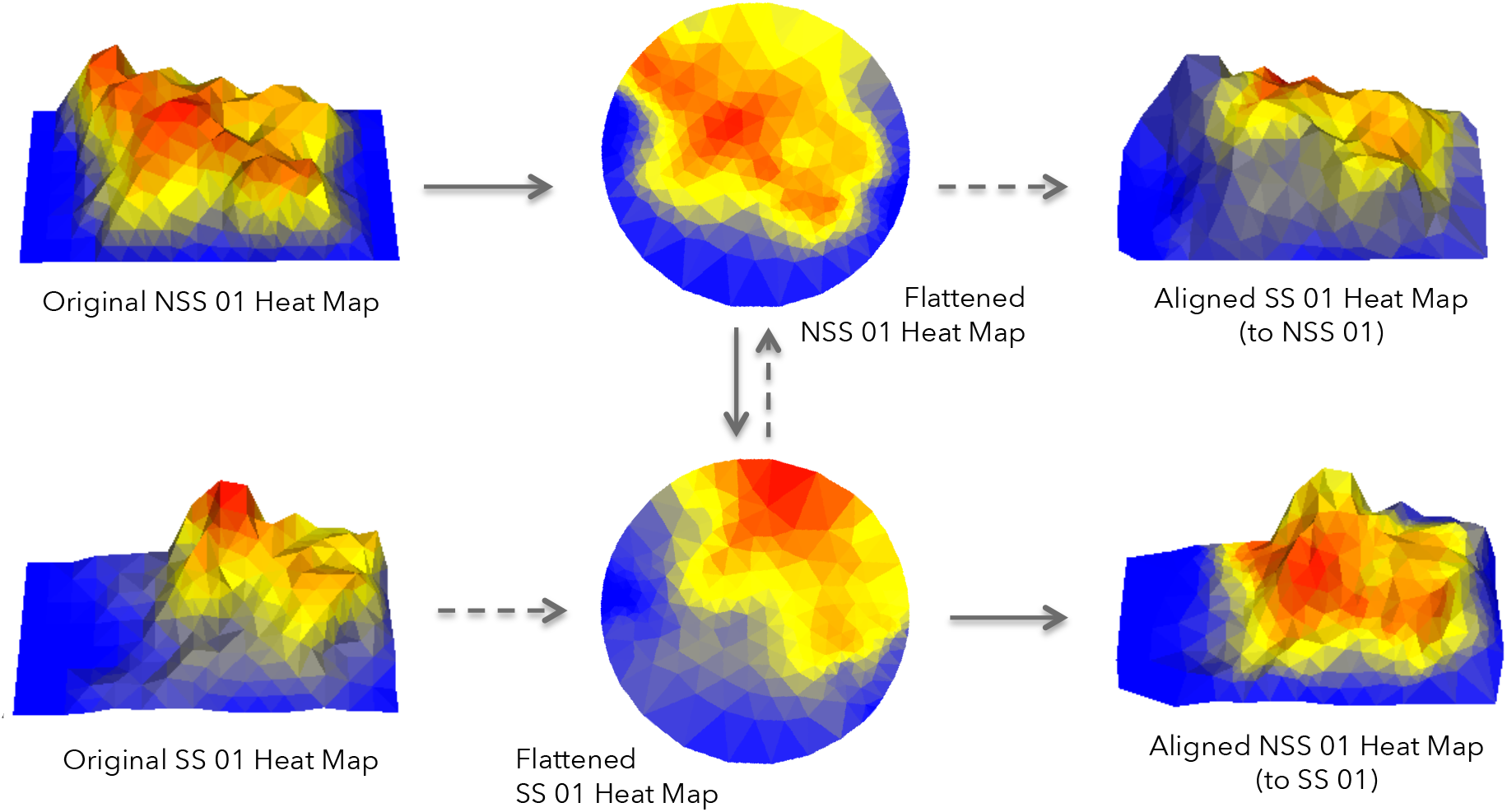
The approximately uniform Delaunay triangulations corresponding to the heat maps of the NSS 01 and SS 01 medaka, their conformal flattenings to the unit disk, the symmetric distortion minimizing Möbius transformation from the unit disk to itself, and corresponding alignments.

We can compare the spatiotemporal distribution functions between two animals by taking an appropriate norm between them, but that will be sensitive to a variety of factors that cannot be controlled, such as the position of an animal when the substance is administered. Mirroring our formulation of BDD, we must build some flexibility into our model by first aligning the corresponding surfaces before calculating a norm between them.

To align a pair of Riemannian surfaces *S*_1_ and *S*_2_, we look at the space of diffeomorphisms between them. A differentiable map *f*: *S*_1_ → *S*_2_ is a *diffeomorphism* if it is a bijection and its inverse *f*^-1^ is also differentiable. A diffeomorphism gives a point-to-point correspondence between surfaces in a way that does not introduce any tears or folds, but it can distort the shape of a surface in unusual ways, for instance by stretching it. Ideally, we would like to find a diffeomorphism that does not distort *S*_1_ while mapping it onto *S*_2_; such a mapping is called an *isometry*. Unfortunately, an isometry between *S*_1_ and *S*_2_ may not exist. Any diffeomorphism between surfaces with different areas, for instance, will have to stretch one of the surfaces somewhere. While we cannot hope to find an isometry between *S*_1_ and *S*_2_, we can always find a map that preserves angles, such a map is called *conformal*. This is the content of a deep result in geometry called the *Uniformization Theorem*.

##### Theorem 6

(Uniformization Theorem, [50]). *Every simply connected Riemann surface is conformally equivalent to one of three Riemann surfaces: the open unit disk, the complex plane, or the Riemann sphere*.

While a conformal mapping *f*: *S*_1_ → *S*_2_ may distort distances, it will stretch *S*_1_ uniformly in all directions at each point. This defines a function λ_*f*_: *S*_1_ → ℝ that maps each point *z* in *S*_1_ to the factor by which vectors at p are stretched by *f*. The map *f* is an isometry if and only if λ_*f*_(*z*) = 1 for all *z* ∈ *S*_1_. Accordingly, one can measure how far a conformal diffeomorphism is from being an isometry be calculating how far the stretching factor is from 1 at each point. [51] introduced the *symmetric distortion energy* of a conformal diffeomorphism *f*: *S*_1_ → *S*_2_ with dilation functions λ_*f*_: *S*_1_ → ℝ and λ_*f*-1_: *S*_2_ → ℝ as

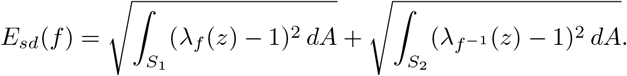

In our context, an optimal alignment between *S*_1_ and *S*_2_ will be a conformal diffeomorphism that minimizes the symmetric distortion energy. We can then measure the difference between two aligned surfaces by calculating the *L*_2_-norm between their corresponding functions.

##### Definition 7.

Let *S*_1_ and *S*_2_ be the surfaces of spatiotemporal distribution functions *ρ*_1_: Ω → ℝ and *ρ*_2_: Ω → ℝ, respectively, and for every diffeomorphism *f*: *S*_1_ → *S*_2_ and *z* ∈ Ω let *z_f_* denote the unique point such that *f*((*z, ρ*_1_(*z*))) = (*z_f_, ρ*_2_(*z_f_*)). The *conformal spatiotemporal distance* between *S*_1_ and *S*_2_ is given by

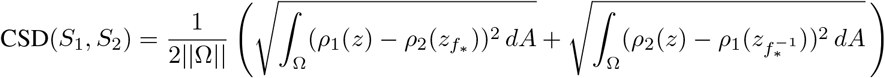

where ║Ω║ denotes the area of Ω and *f*_*_ = argmin_conformal *f*_ *E_sd_*(*f*).

#### 4.3.2 Implementation

To approximate the spatiotemporal distribution function for each fish, we subdivide the rectangular cross-section of the experimental enclosure into an *m* × *n* rectangular grid and assign to each subrectangle the percentage of time spent in that region throughout a specified duration of time. We model the surface corresponding to this function by triangulating each of the rectangular regions and taking the induced triangulation on the corresponding set of points in ℝ^3^. To approximate a conformal mapping from this surface to the unit disk, it will be helpful to have a regular or uniform (all of the edge lengths are approximately equal) triangulation of the surface. Accordingly, we subdivide the edges of our preliminary triangulated surface so that all of the edge lengths are within a specified tolerance of equality and apply the Bowyer-Watson algorithm [52, 53] to obtain a Delaunay triangulation of the resulting vertex set.

We apporixmate the set of conformal mappings between two such triangulations with the Discrete Uniformization theorem [54]. In particular, we use an algorithm developed by [55] to find a discrete conformal mapping 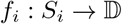 from each triangulated surface / heat map to a triangulation of the unit disk. Once we have these conformal flattening maps, *f*_1_ and *f*_2_, we can express every conformal mapping *f*: *S*_1_ → *S*_2_ as 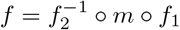 where 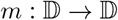 is a *Möbius transformation* of the unit disk. Finding an optimal alignment between *S*_1_ and *S*_2_ is then reduced to finding a Möbius transformation 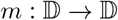 whose composition 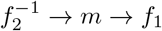 minimizes the symmetric distortion energy. This is illustrated in Figure 7.

A well known application of Schwarz’s Lemma [56] in complex analysis is that every Möbius transformation from D to itself is determined by which point is sent to the origin and a rotation. For stability reasons, we map the centroid of each surface to the origin [51]. This also reduces the minization problem to a single rotation, which we approximate by brute force, although there are algorithms in the literature for this [25]. We use the following discrete analogue of the symmetric distortion energy presented in [51]: If *T*_1_ and *T*_2_ are triangulations of smooth Riemannian surfaces *S*_1_ and *S*_2_, respectively, and *f*: *S*_1_ → *S*_2_, is a conformal mapping, then

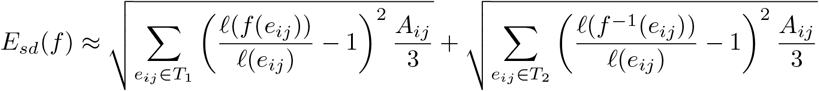

where *e_ij_* denotes an edge in *T*_1_ or *T*_2_, *ℓ* denotes the length of an edge (or its image), and *A_ij_* is the sum of the areas of the two triangles that contain the edge *e_ij_*. Here the sums run over all of the interior edges in *T*_1_ and *T*_2_ respectively.

**Figure 8:**
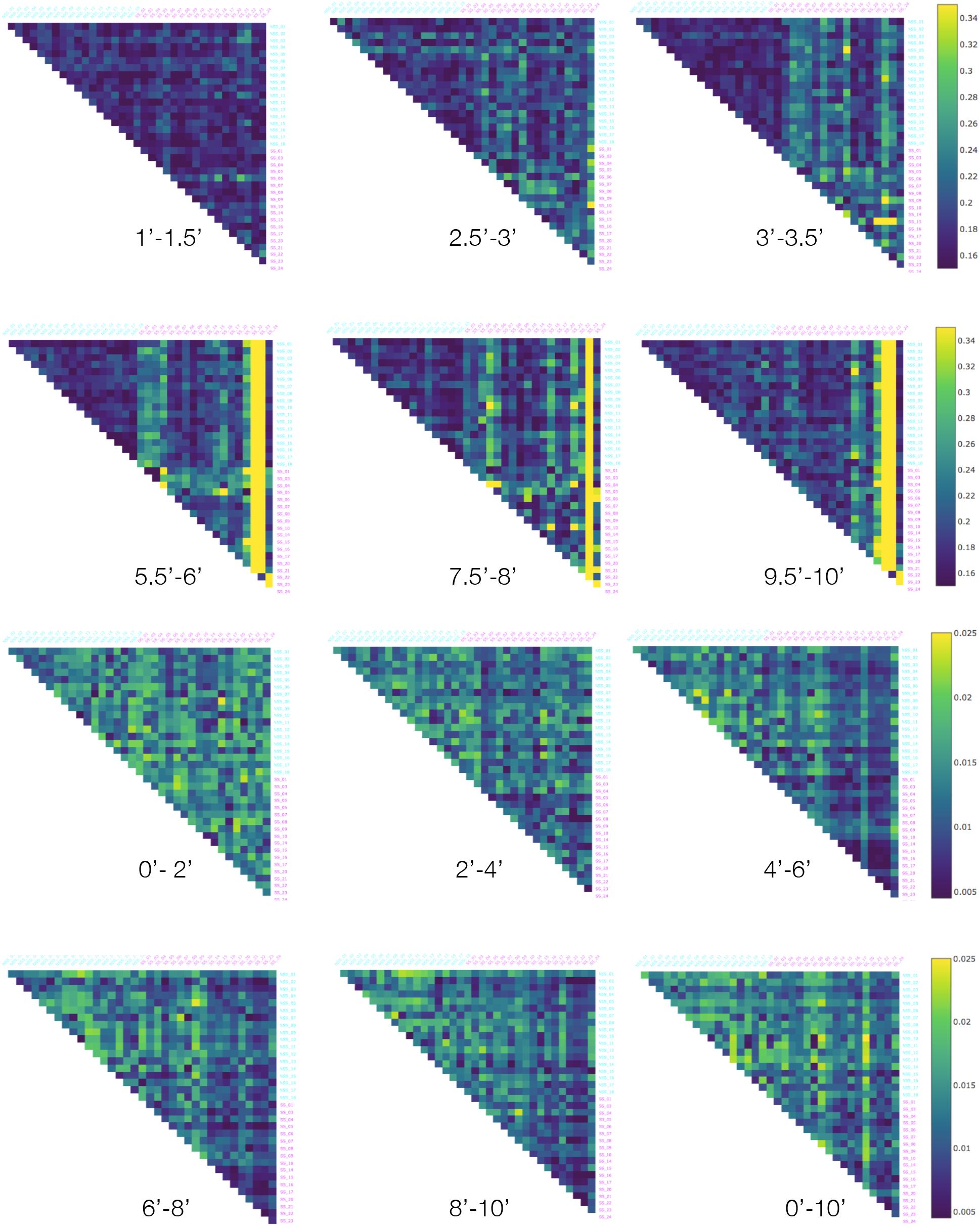
The Behavioral Distortion Distances (upper half) and Conformal Spatiotemporal Distances (lower half) presented in Figures 2 and 3 where the rows and columns are ordered by type rather than centrality.

## 5 Acknowledgments

This work was supported by Yale-NUS College through grants R-607-265-225-121 and IG16-LR003. The authors acknowledge the support from Haroun Chahed, Goh Rui Zhe and Maharshee Karia in the initial stages of the project.

## 6 Author Contributions

ASM and MTS conceived and designed the project, and wrote the manuscript with assistance from SG. ASM provided the data-set to test the methods and interpreted the results. MTS developed the models and conducted the computational experiments. SG developed the software package with assistance from MTS.

### 7 Supplementary Figures & Tables

The following figures and tables were referenced in the Results section.

## Footnotes

2 For each duration *d*, the set of pairs of non-overlapping pairs of intervals can be identified with a triangular region in ℝ^2^ with area (10 – 2*d*)^2^/2).

